# 3D Packing Defects in Lipid Membrane as a Function of Membrane Order

**DOI:** 10.1101/591784

**Authors:** Madhusmita Tripathy, Anand Srivastava

**Affiliations:** Molecular Biophysics Unit, Indian Institute of Science Bangalore, Bangalore 560012, India

**Keywords:** Lipid Packing Defects, Lipid Order, Protein Localization, Membrane-Protein Association

## Abstract

Lipid membrane packing defects are considered as essential parameter that regulates specific membrane binding of several peripheral proteins. In absence of direct experimental characterization, lipid packing defects and their role in the binding of peripheral proteins are generally investigated through computational studies, which have been immensely successful in unraveling the key steps of the membrane-binding process. However, packing defects are calculated using 2-dimensional projections and the crucial information on their depths is generally overlooked. Here we present a simple yet computationally efficient algorithm, which identifies these defects in 3-dimensions. We employ the algorithm to understand the nature of packing defects in flat bilayer membranes exhibiting liquid-ordered (*L_o_*), liquid-disordered (*L_d_*) and co-existing *L_o_*/*L_d_* phases. Our results indicate the presence of shallower and smaller defects in the *L_o_* phase membranes as compared to the defects in *L_d_* and mixed *L_o_*/*L_d_* phase membranes. Such analyses can elucidate the molecular scale mechanisms that drive the preferential localization of certain proteins to either of the liquid phases or their interface. Moreover, on the methodology front, our analyses suggest that the projection based 2-dimensional calculation of packing defects might result in inaccurate quantification of their sizes - a very important feature for membrane association of protein motifs, thus advocating the importance of the 3-dimensional calculations.

## 1 Introduction

Biological membranes do not merely act as selectively permeable boundaries that provide shape and stability to the cell and its organelles, but they also play critical roles in various physiological processes such as transport, signaling and trafficking [1–4]. For most of the cellular functions, role of specific lipids [5–7], membrane electrostatics [8–10] and membrane curvatures [11, 12] are quite well studied with respect to the folding, dynamics and functions of membrane proteins as well as with respect to preferential localization and adsorption of peripheral protein to membrane surface [13, 14]. Another physical feature, which has recently been of great interest in the area of peripheral protein adsorption and protein sorting along the secretory pathways, is the lipid packing defects [15, 16]. Lipid packing defects are identified as those regions on the surface of the membrane where the hydrophobic lipid tails are transiently exposed to the aqueous polar environment. It has been shown that amphipathic motifs of peripheral proteins can sense these packing defects and subsequently bind to the lipid membrane by inserting their bulky hydrophobic residues into them [15–21].

Due to their transient nature and small size, the experimental characterization of lipid packing defects has been elusive and the measurements are carried out indirectly through sensors such as membrane-anchored proteins and amphipathic lipid packing motifs [22, 23]. Several studies have used indirect experimental assays to highlight the role of lipid defects in specific localization and preferential partitioning of peptides and proteins on membrane surface [11, 13, 17, 18, 20, 24–26]. On the other hand, the contribution of computational tools and methods to characterize lipid packing defects has been quite compelling. These tools have proven to be very useful in elucidating the process of binding of amphipathic motifs and their subsequent folding into *α*-helix in presence of packing defects, where the motif folding and defect coalescence has been shown to share a dynamic cooperativity [15–18]. In this line, the scientific contributions from two groups, Voth and co-workers [15, 18] and Antonny and co-workers [16, 17, 19], need a special mention. Voth group was among the first to compute the lipid packing defects structures and distribution as a function of membrane curvature in model atomistic lipid membranes and show the effects of such defects on amphipathic helix (AH) folding into the membrane. They used a solvent accessible surface area (SASA) based method to identify the packing defects and projected the defects to a 2-dimensional plane for subsequent analysis. Antonny’s group, on the other hand, used a 2-dimensional cartesian grid-based method that scanned the lipid surface to identify exposed lipid tail atoms. Based on the nature of the atom encountered, they classified the defect to be either geometric or chemical. Very recently, Antonny and co-workers optimized their algorithm and presented it in the form of a bioinformatic tool, named PackMem [27], where they also characterize packing defects to be shallow or deep. However, these elegant schemes still analyze the lipid packing defects in 2-dimensions (2D) and report them as area though such defects are actually 3-dimensional voids on the lipid membrane surface [28]. Analyzing the defect projection is a technical simplification, where there is a possibility that some crucial information might get lost. The basic reason behind such simplification is the vast number of lipid atoms present in the membrane, which make a 3-dimensional grid search for atom overlap extremely time consuming. A scheme which can efficiently take care of the grid search should be able to fix this issue and identify the packing defects as voids in 3-dimensions (3D).

In this work, we present a simple and robust algorithm to calculate the lipid membrane packing defects in 3D. This algorithm is rooted into free-volume based calculations, which identifies the defect pockets by scanning a local (grid) volume around each atom in the lipid membrane. This is in contrast to the usual grid-based comparison where each atom is compared to every grid point to check for overlaps. The local search, thus, reduces the computational cost. We use this protocol to identify the packing defects in three different ternary lipid membrane systems, DPPC/DOPC/CHOL, PSM/DOPC/CHOL, and PSM/POPC/CHOL, each exhibiting pure *L_o_*, pure *L_d_*, and mixed *L_o_*/*L_d_* phases at various lipid compositions [29–32]. In the last two decades, the nature of packing defects on membranes is well studied with respect to curvature sensing, AH insertion, membrane-association of peripheral proteins and protein/lipid trafficking and localization [15, 17]. In the light of phase gradient in membranes across organelles [28] and the functionally important scale-rich phase organization in plasma membrane [24, 33, 34], it is important to also the explore the nature of packing defects in lipid bilayers as a function of orderliness of the membrane. Recently, we have shown that local topological rearrangements of lipids in liquid ordered regions are distinctly different from lipids found in liquid disordered regions and suggested that these topological rearrangements may depend on the “wiggle space” available to the lipids [31, 32]. In this work, we calculate defects in 3D and find that the distribution of defect pocket sizes turns out to be remarkably different from the projection based 2-dimensional calculation. Our data also provide important insights about defect size and depth distribution in membrane that exhibit purely *L_o_*, purely *L_d_* and co-existing *L_o_*/*L_d_* phase behavior.

The rest of the paper is organized as follows. The details of the atomistic simulation trajectories analyzed in this work are discussed in Section 2, along with a thorough discussion on the proposed algorithm to identify hydrophobic packing defects in lipid membranes. In Section 3, we provide and discuss results on the spatial, size and depth distributions of lipid packing defects for all the nine model lipid membrane systems studied in this work. Subsequently, to rigorously assess the usefulness of the methodology, we discuss the scope and limitations of the algorithm in Section 4. We conclude in Section 5 with our important results and probable future directions.

## 2 Methods

### 2.1 Atomistic Simulation Trajectories of Model Lipid Membranes

The set of atomistic simulation trajectories of lipid membrane system, which are used to benchmark the proposed algorithm, are borrowed from Edward Lyman’s group at the University of Delaware. The set consists of three different ternary lipid systems: DPPC/DOPC/CHOL, PSM/DOPC/CHOL, and PSM/POPC/CHOL, each simulated at three carefully chosen compositions at which the system exhibits pure *L_o_*, pure *L_d_*, and co-existing *L_o_*/*L_d_* phases. The *μs* long simulations were carried out on Anton supercomputer [35, 36], where the system snapshots were collected at every 240 *ps*. Some of the important details of the trajectories are summarized in Table 1. Further details on the technicality of the system set up and simulation protocol can be found in the original references [29, 30].

**Table 1:** Simulation details of the nine atomistic membrane systems analyzed in this work.

| Lipid system | Composition | Phase | Temperature (K) | Total Lipids | Simulation Duration ($\mu s$) |
| --- | --- | --- | --- | --- | --- |
| DPPC/DOPC/CHOL | 0.55/0.15/0.30 | Lo | 298 | 512 | 5.05 |
|  | 0.29/0.60/0.11 | Ld | 298 | 400 | 2.02 |
|  | 0.37/0.36/0.27 | Lo/Ld | 298 | 380 | 9.43 |
| PSM/DOPC/CHOL | 0.64/0.03/0.33 | Lo | 295 | 560 | 0.95 |
|  | 0.15/0.82/0.03 | Ld | 295 | 574 | 0.81 |
|  | 0.43/0.38/0.19 | Lo/Ld | 295 | 452 | 4.54 |
| PSM/POPC/CHOL | 0.61/0.08/0.31 | Lo | 295 | 560 | 9.08 |
|  | 0.23/0.69/0.08 | Ld | 295 | 560 | 1.14 |
|  | 0.47/0.32/0.21 | Lo/Ld | 295 | 506 | 6.88 |

### 2.2 Calculating Hydrophobic Defects

To identify the packing defects on a membrane surface, we follow a grid-based free volume method. The defects are defined as the surface of the hydrocarbon tails of the lipids that are exposed to a probe, and are identified in a two-step calculation. In the first step, we identify the excluded or free volume of the surface of a membrane leaflet and in the second, the defect pockets are located. In the following sections, we discuss these two steps in details.

#### 2.2.1 Identifying Membrane Surface Free Volume

To begin with, we consider one leaflet of the lipid bilayer lying on the XY plane. We define a cutoff Z value and select all the atoms of the monolayer that lie above the cutoff. The Z cutoff is arbitrary and can be efficiently chosen based on a simple visual inspection of the trajectory to be analyzed. A very low cutoff will lead to selection of a large number of atoms and increase the computational cost, while a very high cutoff may miss out important atoms over the trajectory. Therefore, a Z value is chosen which selects sufficient atoms from the monolayer, while making sure that all the lipids lie well above the Z cutoff over the entire trajectory. For simulation trajectories where the origin of the coordinate system is fixed at the center of the bilayer, Z=0 can serve as a decent choice. This selection renders a flat bottom to the membrane monolayer. The monolayer is then mapped to a 3-dimensional box with the flat bottom of the monolayer at the bottom face of the box. Periodic boundary conditions are applied along the X and Y directions, thus fixing the lateral boundaries as per the corresponding simulation box dimensions. The application of periodic boundary condition ensures precise identification of defects that are located near/at the boundaries of the simulation box. The height of the box is chosen to be sufficiently high, so as to keep all the surface atoms well within the box. The coordinates of all the atoms in the box are then shifted such that the whole reference box lies in the positive XYZ quadrant, with origin at the corner of the bottom face. Subsequently, the box is mapped to a 3-dimensional grid of 1 Å resolution, and we identify and remove grid points that lie within one van der Waals (vdW) radius of any atom *plus* a probe radius. We use the vdW radius of atoms as specified by the forcefield and a probe radius of 1.4 Å, which is approximately the radius of a water molecule.

The general protocol for this part of the calculation is to compare the position of each grid point to that of each atom in the reference box to check for overlap. This comparison is usually achieved in two nested loops with either the grid index as the outer loop or the atom index and thus, is computationally expensive. We use a protocol that avoid such expensive nested loops over all atoms/grid points and rather compares an atom to only its neighboring grid points to check for overlapping. The idea is to consider a local 3-dimensional grid around each atom, in the selection, and compare the grid points within this local grid for possible overlap. As the corner of the reference box is kept at the origin and the grid spacing is 1 Å, we can identify the nearest neighboring grid points of any atom quite easily: it is the integer value of the position vector of the atom in any direction. In case the grid spacing was something different that 1 Å, it would be the integer value of the ratio of the position vector and the grid spacing. Thus, considering one atom of the monolayer, we can easily locate the grid points that lie within its vdW radius plus one probe radius in all directions. These grid points constitute a local 3-dimensional grid around the central atom. We now compare the distance between the position of the central atom and all the grid points within the local 3-dimensional grid around it. If this distance is found to be less than its vdW radius *plus* the probe radius, we remove the grid point as it overlaps with the central atom. We repeat the same procedure for all the atoms in the leaflet. The algorithm for this part of the calculation is given below.

~~~
*initialize all grid points (of the entire box) to* 0
**for** *atom i in the box*
   *atom type*[*i*] → *t*
   *radius*[*t*] → *VDW radius*[*t*] + *probe radius*
   *lower boundary of the local box* → (*int*)(*r* − *radius*[*t*])
   *upper boundary of the local box* → (*int*)(*r* + *radius*[*t*])
   *check for overlapping of atom i position and grid points within the local box*
   **if** *grid point is overlaping*
      *grid point value* → 1
   **endif**
**endfor**
~~~

At the end of this analysis, grid points with *grid point value* 0 constitute the free volume of the leaflet selection under consideration, i.e., the excluded volume of all the atoms in the box (selected from one layer of the lipid bilayer). Starting from the rugged surface of the leaflet, this excluded volume extends all the way to the top of the reference box (Fig. S1 in the SI). One can visualize this excluded volume as a mould or cast of the leaflet under consideration. With the excluded volume information available, we next identify the defect pockets on the membrane surface.

#### 2.2.2 Identifying Membrane Defect Pockets

The flat bilayer membranes that are studied in computer simulations are not perfectly flat and exhibit transverse fluctuations that are much smaller in length scales as compared to the membrane’s intrinsic curvature. These fluctuations can play a very important role in the formation of the defect pockets in such flat membranes. We, therefore, look for the defect pockets surrounding each lipid in its local neighborhood.

For this part of the analysis, we define the bottom carbon atom of the glycerol group, from where the two tails bifurcate, of lipids and the hydroxyl group oxygen atom of cholesterols as the reference sites. At this point, it should be noted that in earlier works on hydrophobic defects, the lipid membranes were composed of either a single lipid component or a mixture without cholesterol [15, 16, 19]. As cholesterols lie deeper within the membrane as compared to lipids, neglecting them will introduce inaccuracy in defect pocket calculation. Also, the interaction between water molecules and the lipids can get sufficiently modified in the presence of nearby cholesterols. Therefore, we also consider cholesterols while locating defect pockets. Intuitively, all the reference atoms never fall on a plane and, and so, do the local defects. We define defect pockets as local excluded volume, which we obtain from the earlier calculation, that lie in the neighborhood of the reference sites and extend below them along the Z-direction, thus exposing the hydrophobic tails of the lipids and the aromatic rings of the cholesterols. The neighborhood is defined around the reference sites in the XY plane, within a local 3-dimensional grid of length 4 *times* the probe radius *plus* the vdW radius of the reference atom. With these conditions, the free volume grid points that lie above the membrane surface are automatically discarded and we are left with local defect pockets lying on the membrane.

~~~
*initialize defect grid points to* 0
**for** *reference atom j in the box*
   *atom type*[*j*] → *tt*
   *neighborhood*[*tt*] → *VDW radius*[*tt*] + (4 × *probe radius*)
   *lower boundary of the local box* → (*int*)(*r_ref_* − *neighborhood*[*tt*])
   *upper boundary of the local box* → (*int*)(*r_ref_* + *neighborhood*[*tt*])
   *check for overlapping of reference atom j position and grid points within the local box*
   **if** *grid point lies within neighborhood*[*tt*] *of the reference site in XY plane and below z_ref_*
      *defect grid point value* → 1
   **endif**
**endfor**
~~~

The grid points with *defect grid point value* 1, constitute the defect pockets (Fig. S2 in the SI). The algorithms discussed in this and the last subsections provide the coordinates of grid points as output, which constitute the defect pockets on the leaflet at a given snapshot in the trajectory. This data is collected over the entire trajectory, which can be subsequently used to analyze the defect structure. Figure 1 shows the membrane defects identified using this method in a DPPC/DOPC/CHOL bilayer system exhibiting *L_d_* phase. As highlighted in the figure, the defect pockets can have distinct 3-dimensional geometry and therefore, a simple 2-dimensional projection cannot capture their structure accurately.

**Figure 1:**
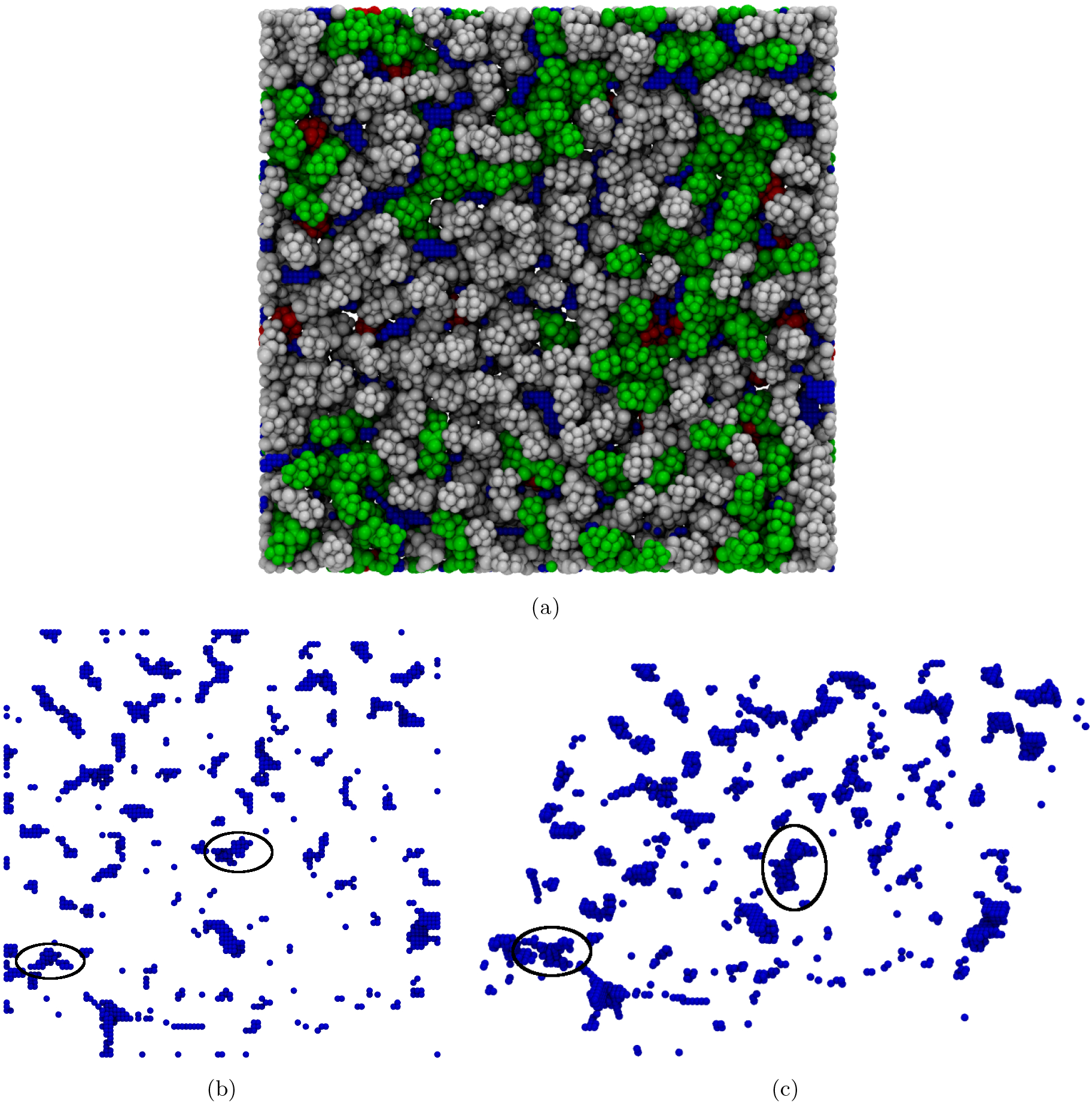
(a) A top view of one leaflet of DPPC/DOPC/CHOL bilayer system, exhibiting *L_d_* phase, showing the membrane packing defects. Lipids and defects are shown using vdW representation: DPPC in green, DOPC in white, cholesterol in red, and defects in blue. Top (b) and side (c) views of the defect grid points. Defects pockets such as those highlighted have distinct 3-dimensional geometry. The figures have been rendered using the visualization tool, VMD [37].

As one can expect, this calculation can (and does, see Fig. S1 in the SI) also identify grid points that lie well within the membrane as defects. One can employ conditions to avoid such occurrence, such as the defect pocket should always originate from the reference atom plane and extend below. However, as the lipid tails are very closely packed, such ambiguous defect pockets are statistically insignificant in number and does not affect the final results.

In Table 2, we present some data on the computational cost of the defect calculation, as proposed in this section. As the lipid systems undergo phase separation over the simulation run, we only choose a short window from each of the equilibrated (already phase separated) atomistic trajectories that constitute our analysis windows. In each case, we choose a suitable Z cutoff based on a simple visual inspection. We identify the defects in each of the membranes over their respective analysis windows following the algorithms provided above and record the runtime. These lipid membrane systems differ considerably in their sizes, with number of lipids varying from 380 to 574 (Table 1). However, in our current implementation of the algorithm, it takes less than half a second to identify defect grid points for a single snapshot of one leaflet for each of the systems. The exact run times are presented Table 2. One should note the run time can be further reduced by choosing the best possible Z cutoff in a flat membrane system. While the algorithm is pretty straightforward, the runtime will always depend on the implementation of the same and thus, a highly optimized version of the code can greatly reduce the computational cost. Lastly, we benchmark the runtime by performing the calculations on a standard desktop, while a powerful computer can lead to substantial reduction in computational cost. We therefore believe that the proposed algorithm can be very useful in analyzing packing defects on large lipid membrane systems and provide useful insight to its biologically significant functions such as membrane-protein association.

**Table 2:** Computation cost (runtime) for identifying the defects in the nine atomistic membrane systems.

| Lipid system | Phase | Analysis Window (snapshots) | Total Runtime (s) | Runtime Per-snapshot (s) |
| --- | --- | --- | --- | --- |
| DPPC/DOPC/CHOL | Lo | 500 | 110 | 0.220 |
|  | Ld | 500 | 96 | 0.192 |
|  | Lo/Ld | 500 | 75 | 0.150 |
| PSM/DOPC/CHOL | Lo | 193 | 38 | 0.197 |
|  | Ld | 165 | 45 | 0.273 |
|  | Lo/Ld | 313 | 56 | 0.180 |
| PSM/POPC/CHOL | Lo | 279 | 57 | 0.205 |
|  | Ld | 501 | 178 | 0.360 |
|  | Lo/Ld | 285 | 66 | 0.232 |

## 3 Results and Discussion

### 3.1 *L_d_* Regions of the Membrane Exhibit Denser Distribution of Packing Defects than the *L_o_* ones

To understand the spatial distribution of defects in lipid membranes exhibiting ordered and disordered phases, we analyze their spatial maps, which were generated in the following steps. The defect pockets were first projected onto the XY plane of the grid. Ten such projections were collected from ten consecutive system snapshots. Subsequently, the set of projected defects were binned in XY plane with bin size 1 Å (the original grid spacing) and averaged over the number of snapshots. Thus, the colorbar in the map indicates the probability that a grid point is a part of a defect. A high probability, therefore, indicates the persistence of a defect pocket in the XY plane, i.e., a localized defect. It should be noted that the system snapshots, considered here, are collected 240 *ps* apart in time. Therefore, considering a large number of snapshots will be pointless as the defects are highly transient structures.

The ten steps averaged spatial maps of defects, calculated for the pure *L_o_* and pure *L_d_* phase membranes of the DPPC/DOPC/CHOL systems, are shown in Fig. 2. Also shown are the top view (in the XY plane) of the corresponding membrane configurations, generated at the middle (the sixth) snapshots of the chosen sets. The relatively straight configuration of the lipid tails in the *L_o_* phase leads to the compact ordering of the lipids upon aggregation. Therefore, the *L_o_* phase system is found to be scarce in defect pockets: a direct consequence of the compact arrangement of the lipids therein, which does not allow wiggle space between the lipids such that the hydrophobic tails can get exposed. On the other hand, the *L_d_* phase membrane displays a high density of defects. Also, given that the spatial maps are averaged over ten consecutive snapshots, one cannot fail to notice that the defects in the *L_d_* system have extended over the 2.4 *ns* window, where as those in the *L_o_* system seem to be localized, exhibiting high probability of defect occurrence.

**Figure 2:**
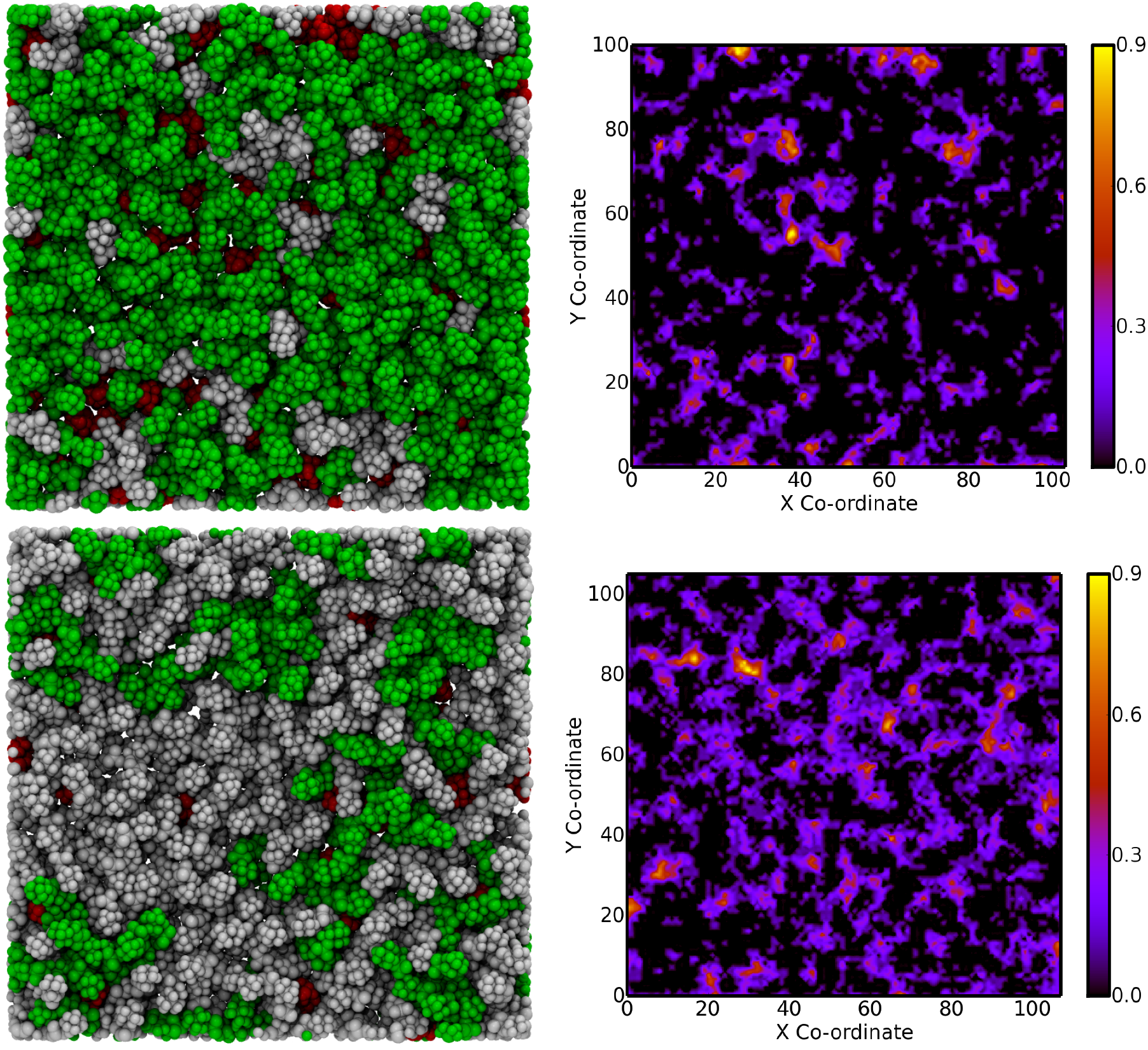
Membrane configurations (left panel) for DPPC/DOPC/CHOL systems exhibiting *L_o_* (top) and *L_d_* (bottom) phases. All lipids are shown in vdW representation with DPPC in green, DOPC in white, and cholesterols in red. Right panel shows the spatial distribution maps of defects therein, with colorbars indicating the defect distribution count or occurrence probability over an interval of 10 snapshots. The axes shown are in A units.

Given the distinct nature of defects in pure *L_o_* and *L_d_* systems, it is intriguing to investigate the same in mixed phase *L_o_*/*L_d_* systems. The results from the three mixed phase systems, considered in this work, is presented in Figure 3. The *L_o_* regions in all these mixed phase membranes are rich in saturated lipids, DPPC in case of DPPC/DOPC/CHOL system and PSM in case of PSM/DOPC/CHOL and PSM/POPC/CHOL systems. As expected, the *L_o_* regions of the membranes are found to be scarce in defect pockets, as in the pure *L_o_* phase membrane. (The small patch on the bottom right in case of DPPC/DOPC/CHOl, right half in PSM/DOPC/CHOl, and near both boundaries along the X-axis in PSM/POPC/CHOL system in Figure 3 left panel.) Similarly, the *L_d_* regions display high density of defects. As in the case of pure *L_o_* system, even the largest defects in the *L_o_* regions stay localized over the ten snapshots (2.4 *ns*): the large defects on top right corner in PSM/DOPC/CHOL system and those on the top left corner in PSM/POPC/CHOL system are few examples of such defects. Thus, while highly localized hydrophobic defects are associated with the ordered regions, the disorderedness of lipids in the *L_d_* regions leads to a high density of defects and make them distinctly mobile. It is therefore evident that our algorithm is able to capture the distinct nature of defects in the ordered and disordered regions of a lipid membrane, which can be useful in understanding the preferential membrane-protein association at distinct regions of the membrane or at their interface [20, 24, 38, 39].

**Figure 3:**
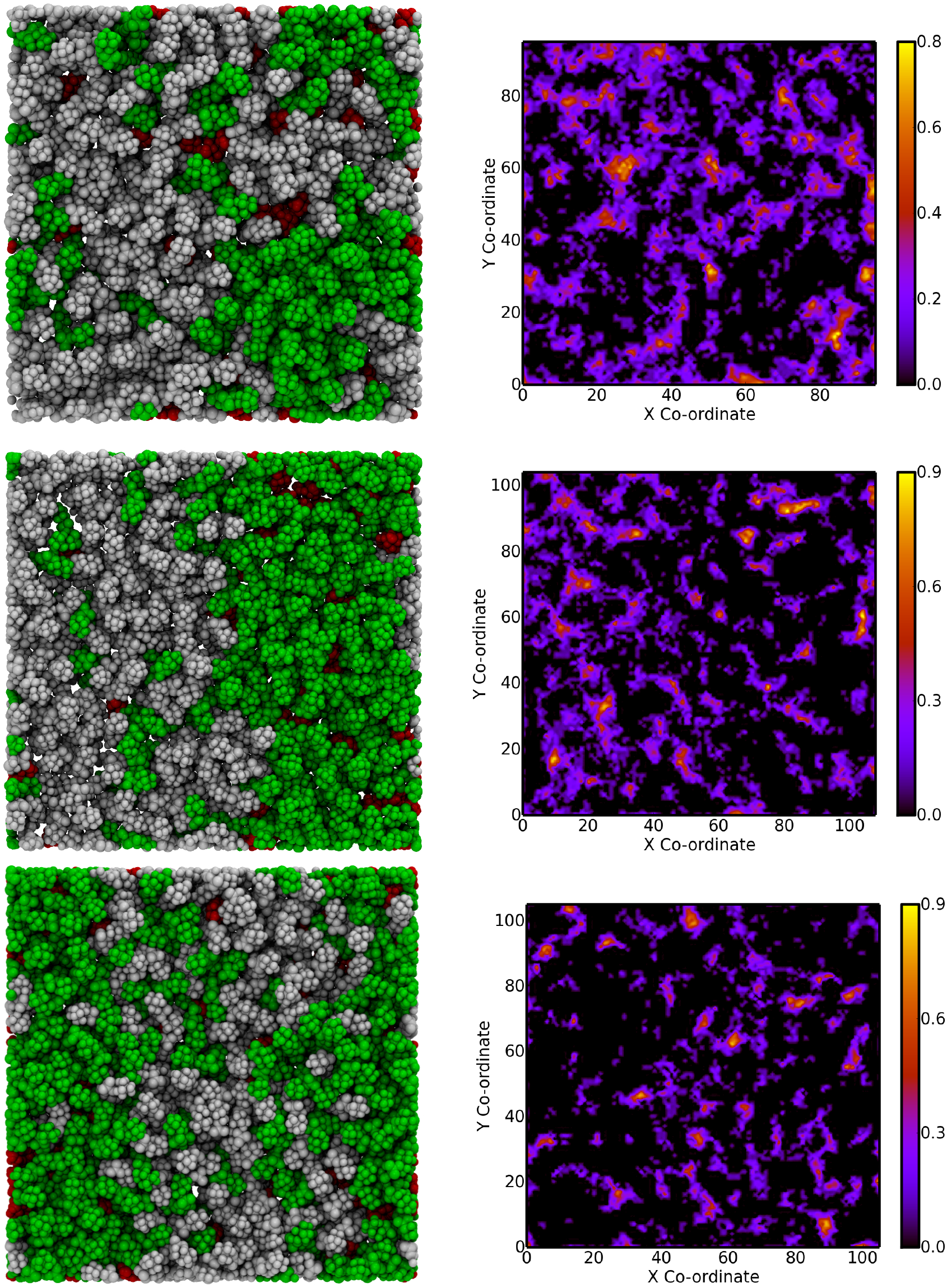
Membrane configurations (left panel) for DPPC/DOPC/CHOL (top), PSM/DOPC/CHOL (middle), and PSM/POPC/CHOL (bottom) systems, each exhibiting mixed *L_o_*/*L_d_* phases. All lipids are shown in vdW representation with saturated lipids (DPPC and PSM) in green, unsaturated ones (DOPC and POPC) in white, and cholesterols in red. Left panel shows the spatial distribution maps of defects over an interval of 10 snapshots. The axes shown are in Å units.

In our earlier studies [31, 32], we observed that the compact topological rearrangement of lipids in the *L_o_* regions make them dynamically steady, which should be the reason for the localization of the defects therein. On the other hand, comparatively faster evolution of lipids in the *L_d_* phase can make the defects more dynamic. We are currently working on establishing any such possible correlation between the evolution of lipids in the membrane and the defect pockets therein, which will be discussed in our future publication.

### 3.2 Size Distribution of Packing Defects Differ Substantially when Calculated in 2D vs. 3D

The most general way to characterize the packing defects is to analyze the distribution of their sizes. The existing protocols to analyze packing defects, which deal with the projection of the defects on a grid plane, represent the defect size as defect area, which is the area of the square grids identified as part of defects. However, here we extract the packing defects as 3-dimensional voids, and thus the calculation of defect size becomes straightforward. We represent these voids as collection of spheres of diameter 1 Å (positioned at each grid point) and calculate defect size as the total number of grid points within it. We count this number by using a simple distance based clustering algorithm. where the grid points, identified from the *defect grid* data (obtained from the algorithms discussed in the last section), which lie within a diagonal distance of each other are merged as a single cluster, and thus form a defect pocket. Mathematically, the exact volume of the void and the size, calculated this way, differs only by a constant factor and therefore, the statistics should be comparable.

In defect size distribution plots, the data points corresponding to very small defects, such as those less than 20, can be conveniently ignored as such small defects appear in all lipid membrane systems irrespective of their nature. On the other hand, very large defects are rare and thus have very low probability of occurrence. Therefore, only the defects with size in the intermediate range are statistically relevant. In general, the defect size distribution, in this range, is found to be exponential and so, the slope of the distribution, plotted in a semi-log scale, indicates the average defect size.

To understand the nature of packing defects in flat lipid membranes, exhibiting pure and mixed liquid phases, we calculate the distribution of their sizes. We first compare these distributions to those obtained from a 2-dimensional defect calculation. To calculate the defect size distribution in 2-dimension, we project the 3-dimensional defects onto the XY-plane and calculate their sizes using our clustering algorithm. In Fig. 4, we compare these distributions for all the nine systems studied in this work. As evident, the defect size distributions, in the range of interest, differ substantially for the two cases and subsequently, the estimated value of average defect size. To quantify this difference, we fit the size distribution data to an exponential function, whose decay constant estimates the average size of the defects in the system and can be found from the slope of the semi-log plot. For DPPC/DOPC/CHOL *L_o_*/*L_d_*, PSM/DOPC/CHOL *L_d_*, and PSM/POPC/CHOL *L_o_* systems, these straight line fits are shown with bold black lines in Fig. 4. The lengths of the straight lines indicate the range over which the data has been fitted. The inverse of the slope estimates the average defect sizes to be 14.1, 14.3, and 7.7 in 2-dimension and 30.3, 29.4, and 14.5 in 3-dimension for the above systems, respectively. The average defect size calculated in 3-dimension is, thus, nearly double of that calculated in 2-dimension (size in our calculations indicates the number of grid points constituting a single defect pocket). We, therefore, emphasize that studies intending to rigorously quantify lipid packing defects should identify them in 3-dimension rather than analyzing a projection of the same.

**Figure 4:**
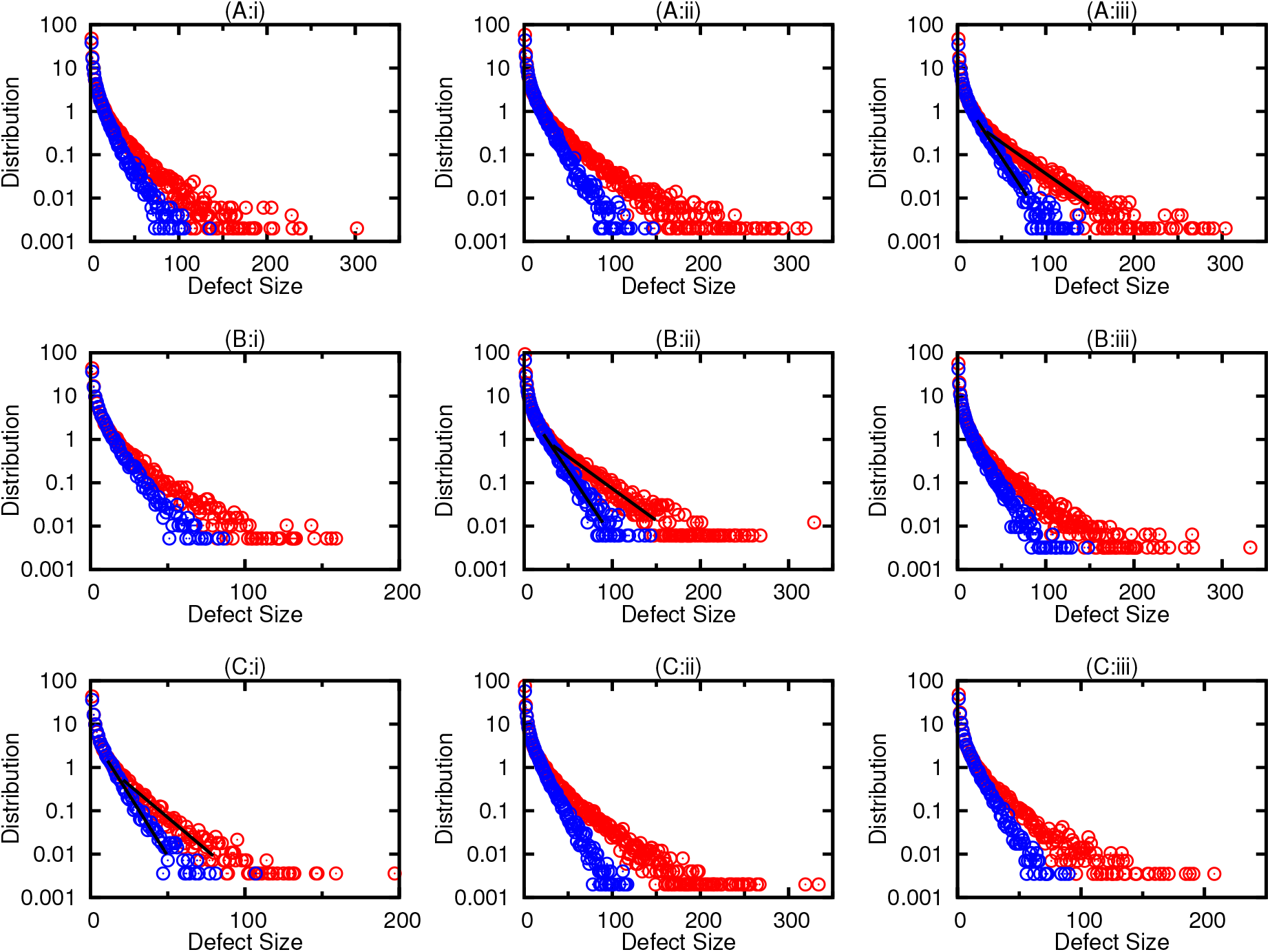
Distribution of defect pocket sizes in 3-dimension (red) and 2-dimension(blue) for (A) DPPC/DOPC/CHOL, (B) PSM/DOPC/CHOL, and (C) PSM/POPC/CHOL systems exhibiting pure *L_o_* (i), pure *L_d_* (ii), and mixed *L_o_*/*L_d_* (iii) phases. Defect size is calculated as the total number of grid points in a defect pocket.

To access the nature of the packing defects across the three lipid mixtures under consideration, we compare their size distributions in Fig 5. In all cases, the *L_o_* phase is found to comprise smaller defect pockets as compared to the *L_d_* phase. This observation can be attributed to the compact arrangement of lipids in an *L_o_* phase that does not allow significant amount of voids to persist on the membrane. The mixed phase exhibit characteristics of both the ordered and disordered phases and therefore, the *L_o_*/*L_d_* phase in PSM/DOPC/CHOL systems is found to exhibit defect sizes that are intermediate to both the pure phases. However, the defect size distribution in the mixed phase DPPC/DOPC/CHOL membrane is found to be similar to that of the pure *L_d_* phase, while that in PSM/POPC/CHOL membrane is similar to that of the pure *L_o_* phase. Therefore, we believe that the mixed phase in DPPC/DOPC/CHOL system is dominated by the *L_d_* subphase, while that in the PSM/POPC/CHOL system is dominated by the corresponding *L_o_* subphase. The spatial distribution of defects shown in Fig. 3 also indicates the same behavior. However, rigorous analyses will be necessary to support this claim, which is the subject of our ongoing investigation.

**Figure 5:**
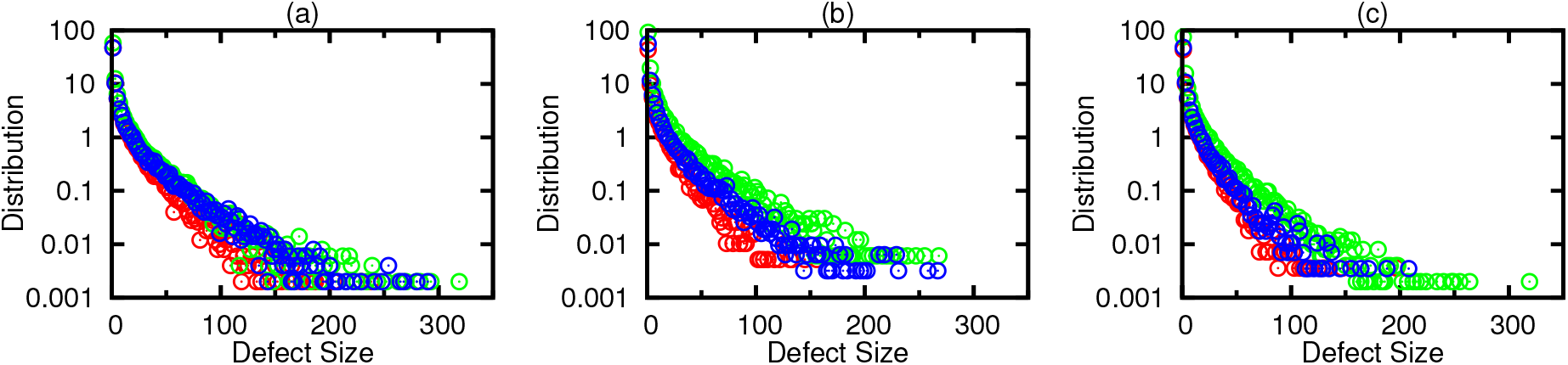
Distribution of defect pocket sizes for (a) DPPC/DOPC/CHOL, (b) PSM/DOPC/CHOL, and (c) PSM/POPC/CHOL systems exhibiting pure *L_o_* (red), pure *L_d_* (green), and mixed *L_o_*/*L_d_* (blue) phases. Every alternate data point is plotted for visual clarity.

### 3.3 *L_d_* and Mixed Phase Membranes Exhibit Deeper Packing Defects than *L_o_* Ones

To further explore the importance of analyzing packing defects in 3D, we calculate the distribution of their depths. The depth of a defect is defined as its net extension along the Z-direction, which is again straightforward to calculate from the *defect grid* data. In Fig. 6, we present these distributions for all the nine systems under consideration.

**Figure 6:**
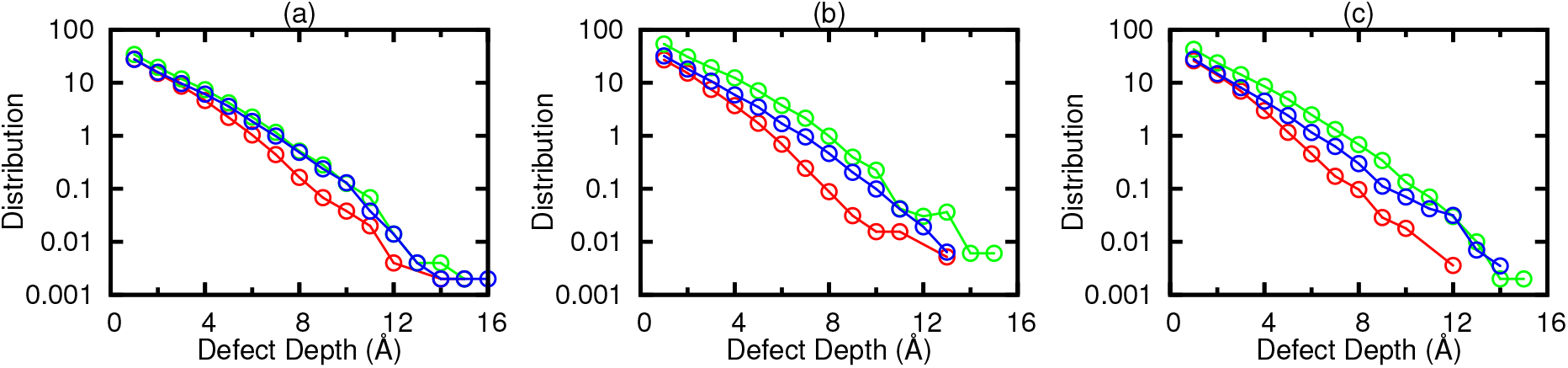
Distribution of defect pocket depths for (a) DPPC/DOPC/CHOL, (b) PSM/DOPC/CHOL, and (c) PSM/POPC/CHOL systems exhibiting pure *L_o_* (red), pure *L_d_* (green), and mixed *L_o_*/*L_d_* (blue) phases. Lines are guide to the eyes.

As earlier, we ignore very shallow (< 6 Å in depth) and very deep (> 12 Å in depth) defects, and focus on the defects in the intermediate range of depths. For all the three lipid mixtures, the membrane exhibiting *L_o_* phase comprise shallower defects than those in *L_d_* and mixed phase *L_o_*/*L_d_* membranes. For example, in PSM/DOPC/CHOL and PSM/POPC/CHOL systems, the defect pockets with depth between 8 and 10 Å differ close to one order of magnitude in their distributions (occurrence probability) between the *L_o_* and *L_d_* cases. In DPPC/DOPC/CHOL systems, this difference is somewhat smaller, but nonetheless pronounced. The defects pockets in mixed *L_o_*/*L_d_* phase membrane are found to be comparable in depth to those in *L_d_* phase membrane for DPPC/DOPC/CHOL systems, while for other two cases, mixed phase membranes exhibit shallower defect pockets as compared to disordered membranes.

The difference in defect depths across the *L_o_, L_d_* and *L_o_*/*L_d_* systems is again as expected. In our recent study [31, 32], we have observed the spatio-temporal evolution of lipids in the ordered and disordered regions of the membrane to be significantly different. This, along with the compact packing of lipids in the ordered regions, does not allow deeper defects to persist. On the other hand, fast moving disordered lipids in the *L_d_* phases leads to larger and deeper defect pockets. Such information on depth distribution of membrane packing defects can be immensely important in the study of membrane-protein association [15–18], where deeper defects can facilitate protein anchoring and subsequent binding. It is needless to mention that such differences in terms of defect depths cannot be identified in a 2-dimensional projection based defect calculation (also see Fig. S3 in the SI). Though the defect analysis tool from Antonny and co-workers [27], PackMem, can identify defects to be shallow or deep, it cannot provide the exact measure of their depths.

## 4 Scope and Limitations

The algorithm proposed in this work, while efficiently handles the job of overlap check, does not impose any fixed parameters. Rather, users can choose various parameters as per their requirement. Firstly, by suitably choosing the Z-cutoff, the number of atoms in the leaflet under consideration can be greatly reduced, thus reducing the computation time. Secondly, for membranes with various species of biomolecules other than lipids, such as sterols, peptides, and proteins, one can choose the reference sites based on the nature of these biomolecules, so as to correctly identify the packing defects around them. As discussed earlier, neglecting such molecules from the calculation can introduce inaccuracy in the defect pocket identification. Lastly, one also has the flexibility of choosing the probe radius and thus identify packing defects of specific nature. In our calculations, we have used a probe radius of 1.4 Å, the approximate radius of a water molecule, so that the defects are the hydrophobic surface available to water. One can also choose the same to be the radius of a bulky hydrophobic residue on a peptide (approximately 3 Å), as used in earlier analyses by other groups [15, 16].

The lipid membranes considered in our analyses are mostly flat and so, both leaflets are essentially identical in nature, in terms of curvature and composition. While we have analyzed the defects in only one of the leaflets, the same analyses can be performed on the other as well with a suitable choice of Z-cutoff. In fact, it is recommended to perform the analyses on both leaflets of the bilayer membrane, so as to improve the statistics of the distribution. Being a benchmarking exercise, we analyze defects in only one leaflet of the bilayer for all the systems, so the comparison is unbiased.

For curved membranes, where lipid packing can be drastically different between the inner and outer leaflets, the characteristics of defects on the two leaflets can be very distinct. For such curved membranes, a Z-cutoff cannot be used to select a subset of atoms from one of the leaflets. In stead, all atoms from one leaflet have to be chosen for analysis and the leaflet can be suitably moved to lie at the bottom boundary of the reference box. It should be mentioned, at this point, that for such a curved membrane, the first part of the proposed algorithm will identify excluded volume that will lie beneath the leaflet itself and so, can be identified as defects in the second part of the calculation. Therefore, to handle such ambiguity, we can impose additional constraints, such as, we can locate carbon atoms near the end of each lipid tail and remove all grid points below it. However, the proposed protocol, of checking overlapping between an atom and its neighboring grid points lying within a local 3-dimensional box, can be used to efficiently identify defect pockets in membranes exhibiting various state of curvature.

## 5 Conclusions

In this work, we provide a coherent framework to systematically characterize lipid membrane packing defects in 3D. The protocol allows the user to choose a subset of atoms from a monolayer to analyze surface defect pockets and the algorithm follows a local grid search method to identify overlaps, thus making the framework computationally very efficient. For all the nine systems considered in this work, summarized in Table 1, it takes less than 1 second to identify defect pockets for a given snapshot of the trajectory. The algorithm takes care of periodic boundary conditions in the lateral directions, such that the defect pockets lying at and near the boundaries are correctly identified. We show that the proposed algorithm yields distinctly different distributions of defect sizes as compared to those from 2-dimensional projection based defect size calculation, which cannot be ignored while providing defect size and population dependent rationale behind biologically important processes. The defect depth information from the 3-dimensional structure of the defect pockets can be crucial while studying the association of peripheral proteins with membrane and while studying selective protein localization as a function of membrane order. The framework is best suited for a flat lipid membrane. However, the same can be extended to curved membranes with a little additional cost of computation. Finally, the algorithm is very simple and thus allows modifications based on user requirements.

We used this method to identify defect pockets in 3 systems of flat lipid bilayers: DPPC/DOPC/CHOL, PSM/DOPC/CHOL, and PSM/POPC/CHOL, each exhibiting pure *L_o_*, pure *L_d_*, and mixed *L_o_*/*L_d_* phases. The *L_o_* phase membranes were found to exhibit smaller and shallower defects than those in the *L_d_* and *L_o_*/*L_d_* phases, which can be attributed to the compact arrangement of lipids in a *L_o_* system. The mixed *L_o_*/*L_d_* phases were found to exhibit defects that are intermediate in size and depth to both the pure phase membranes. The data from the defect calculation are thus in line with the current understanding of membranes exhibiting pure and mixed liquid phases [29–32], and validates the proposed framework. We are currently in the process of mapping out the connection between the nature of defect pockets in such membranes and the spatio-temporal evolution of the membrane lipids. Such a connection can shed light on the molecular origin of the spatial heterogeneities observed in biological membranes that act as platforms for peripheral proteins to bind.

## 6 Acknowledgement

The authors thank Edward Lyman (University of Delaware, USA) for sharing the simulation trajectories and acknowledge Anton supercomputer facility for making the trajectories available. The financial support from Indian Institute of Science Bangalore are greatly acknowledged. AS thanks the startup grant provided by the Ministry of Human Resource Development, India and MT acknowledges the research fellowship from the Department of Biotechnology, India.

## Supplementary Information

**Figure S1:**
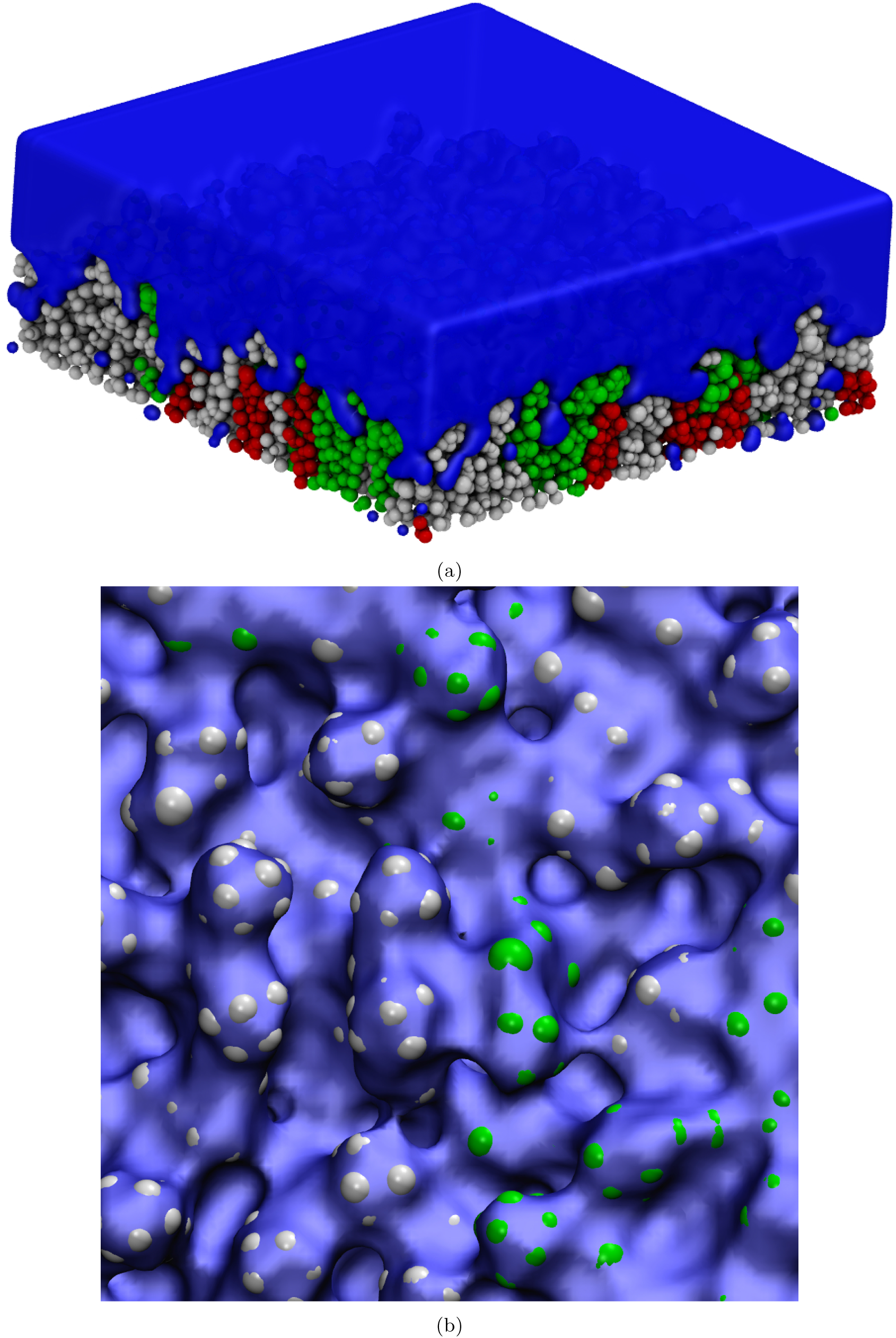
(a) A top view of the membrane surface free volume, calculated using the first part of the algorithm for one leaflet of DPPC/DOPC/CHOL bilayer system exhibiting *L_o_*/*L_d_* phase. The figures have been rendered using the visualization tool, VMD [1]: lipids are shown using vdW representation (DPPC in green, DOPC in white, cholesterol in red) and the free volume is shown using quicksurf representation (in translucent blue). (b) Details of a patch of the leaflet, where blue surface indicates the bottom surface of the free volume that complements the rugged surface of the lipid leaflet. The free volume surface and vdW spheres are not to scale.

**Figure S2:**
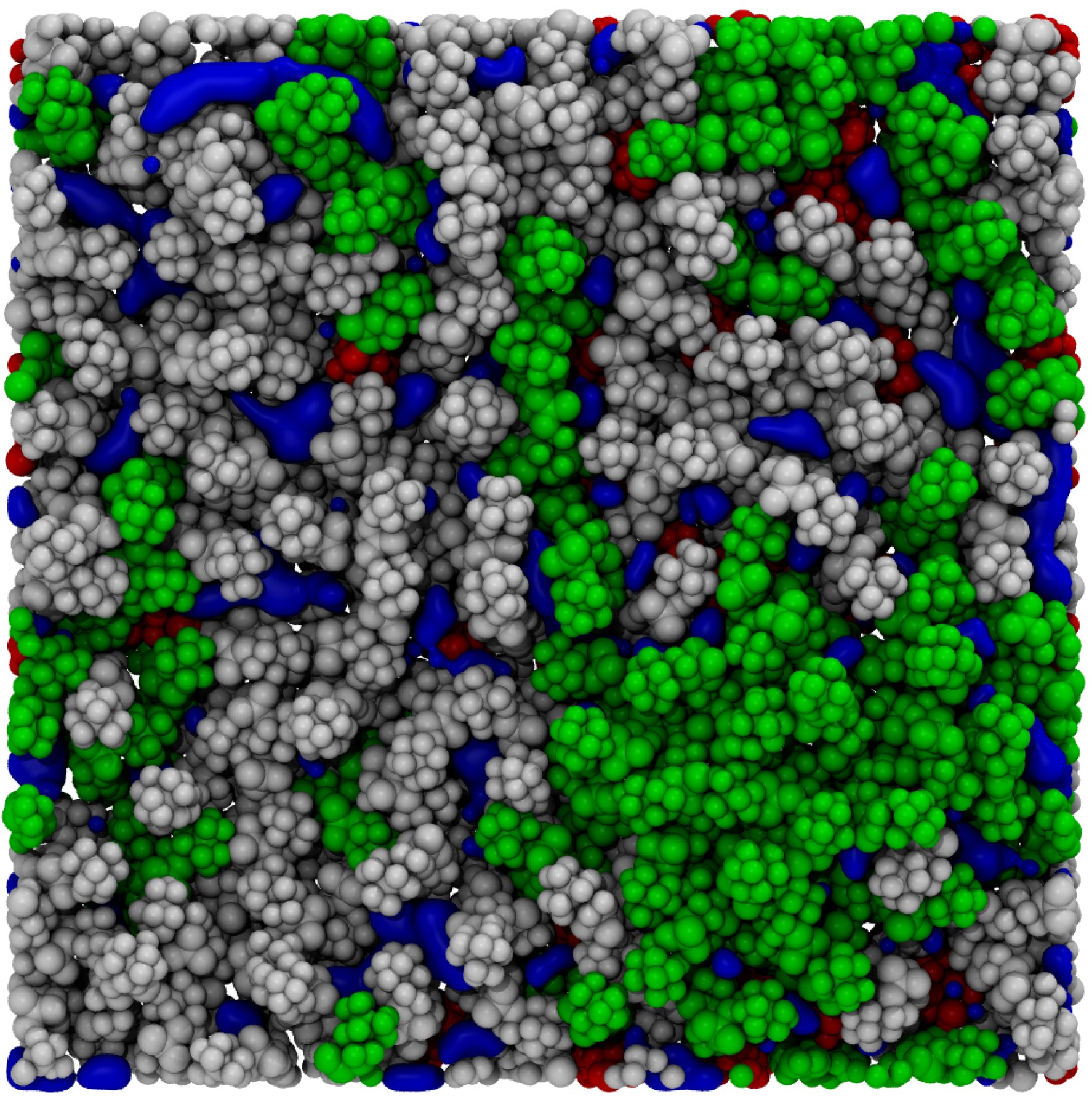
Top view of the DPPC/DOPC/CHOL membrane leaflet, showing the membrane packing defects identified using the second part of the algorithm. The details of the rendition are same as those in Fig. S1.

### Size Depth Ratio of Defects

Assuming a symmetric shape for the defects, a defect of depth *L* will have a size *L*^3^ in 3-dimensions. Thus, the ratio of defect size to the cube of its depth can be a measure of how far the defects are from being symmetric. A value less than 1 will indicate an elongated shape of the defect, while a value more than 1 will indicate a flatter one. In Fig. S3, we plot the distribution of this ratio, calculated for the three systems for which average defect sizes have been calculated in Fig. 4 of the main text: DPPC/DOPC/CHOL, PSM/DOPC/CHOL, and PSM/POPC/CHOL systems exhibiting mixed *L_o_*/*L_d_*, pure *L_d_*, and pure *L_o_* phases, respectively. As evident, a large fraction of the defects have elongated shapes in all the three systems. Consequently, there is disparity in the distributions of defect sizes calculated in 2D and 3D, as shown in Fig. 4 of the main text. While the fraction of elongated defects is highest for the *L_d_* phase, it is the lowest for *L_o_*, in line with the results in Fig. 6 of the main text.

**Figure S3:**
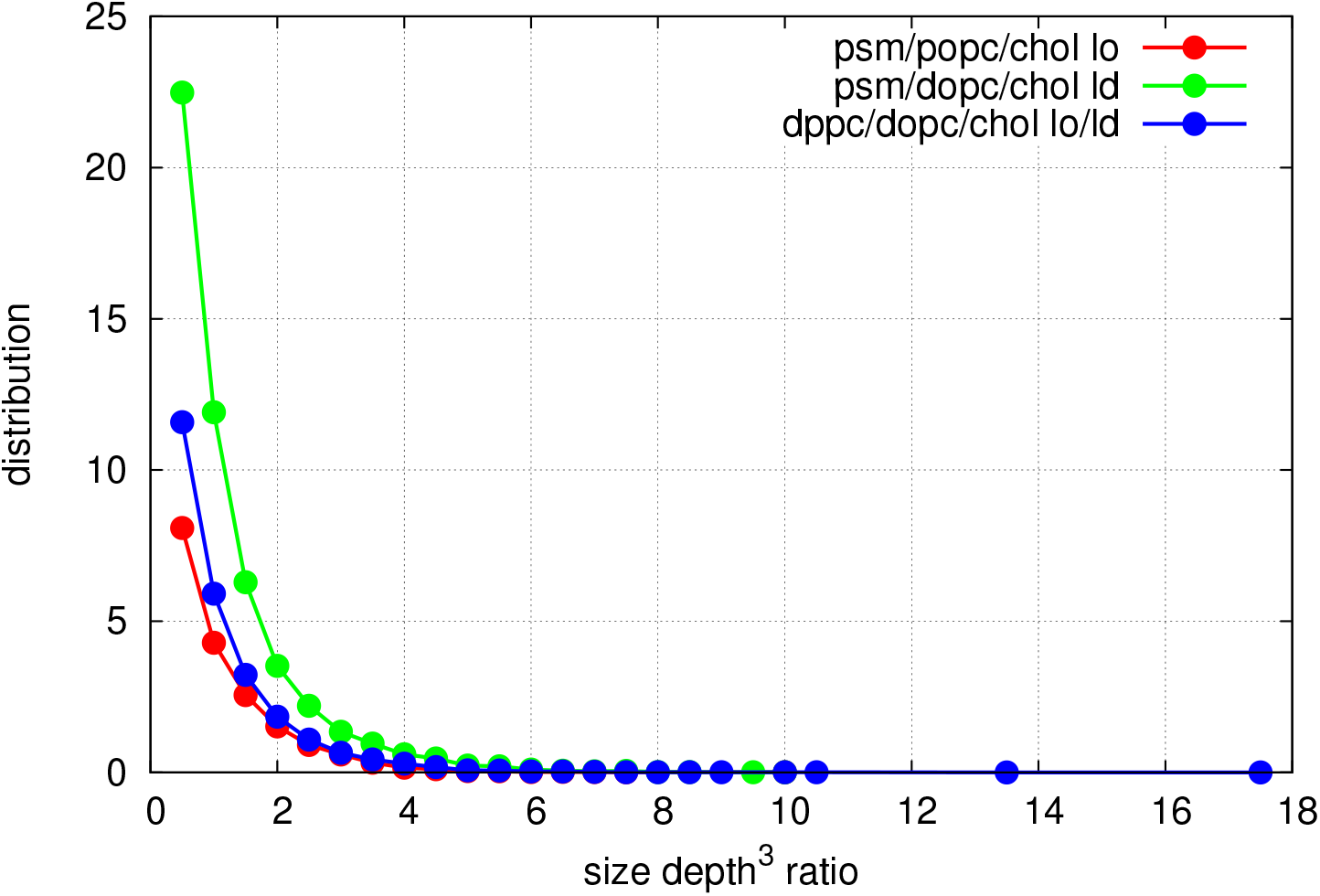
Defect size vs depth cube ratio for DPPC/DOPC/CHOL, PSM/DOPC/CHOL, and PSM/POPC/CHOL systems exhibiting mixed *L_o_*/*L_d_*, pure *L_d_*, and pure *L_o_* phases, respectively. A value ≤ 1 indicates elongated defects while those > 1 indicates flatter defects.

